# Near-critical brain dynamics track effortlessness during meditation

**DOI:** 10.64898/2026.08.07.743470

**Authors:** Evan Lewis-Healey, Morten Kringelbach, Andres Canales-Johnson, Ruben Laukkonen

## Abstract

Effortful cognition is typically associated with controlled, task-constrained neural processing, whereas effortless awareness may require a more flexible, internally driven mode of brain organization. Critical brain dynamics provide a principled framework for characterizing this shift, as systems near criticality are thought to balance stability and flexibility, allowing efficient information processing without excessive control. Transcendental Meditation (TM), characterized by a shift from effortful mental engagement to effortless awareness, offers a natural model for testing this possibility. Here, we investigated whether critical brain dynamics track TM as a global meditative state, or instead reflect moment-to-moment fluctuations in subjective effortlessness. We combined high-density electroencephalography (EEG) with time-resolved phenomenological reports using Temporal Experience Tracing (TET). Experienced TM practitioners (N = 33) and matched controls (N = 33) completed resting-state recordings before and after a 30-minute TM or silent counting control task. Long-range temporal correlations (LRTCs) were quantified using detrended fluctuation analysis, while functional excitation/inhibition (fEI) balance was used to estimate directional deviations from criticality. State-based analyses showed that TM increased alpha and beta LRTCs relative to pre- and post-resting state within meditators, but revealed no robust between-group differences in either LRTCs or fEI balance. In contrast, neurophenomenological analyses showed that subjective effortlessness was robustly associated with increased theta, alpha, beta, and broadband LRTCs, with significantly stronger relationships in meditators than controls. Restricting analyses to low-effort periods further revealed higher beta LRTCs in meditators, a difference missed by conventional state comparisons. These findings identify scale-free neural dynamics as a candidate marker of “letting go” during meditation.

## INTRODUCTION

Meditation is an umbrella term for a group of practices that involve the modulation of attention, often with the aim to cultivate increased mindfulness and wellbeing (Lutz et al., 2008). The variety of practices is extremely vast; different practices entail distinct phenomenological consequences (Jachs, 2021; Laukkonen and Slagter, 2021; Lutz et al., 2015), with extremely profound effects at both the phenomenological (Chowdhury et al., 2023; Laukkonen et al., 2023) and neural levels (van Lutterveld et al., 2025; Potash et al., 2025; Yang et al., 2024) that accompany thousands of hours of practice.

One such meditative discipline is Transcendental Meditation (TM), a standardised mantra-based practice usually conducted twice daily with eyes-closed in 20-minute increments. TM involves the silent repetition of a nonsemantic sound (i.e., “mantra”), as a vehicle to pure awareness (Metzinger, 2020), a state marked by extreme positive valence, coupled with significantly reduced sensory and cognitive content and the preservation of wakefulness (Chandia-Jorquera et al., 2026; Travis, 2014; Travis and Pearson, 2000). One of the fundamental components of the practice is an effortless attention towards the mantra (Mahone et al., 2018), which supports this transcendence into pure awareness states. While there have been a variety of studies highlighting the neural effects of TM (Chandia-Jorquera et al., 2026; Mahone et al., 2018; Travis and Parim, 2017; Travis and Wallace, 1999), there have been no studies to-date that have investigated how critical dynamics within the brain changes as a function of TM.

Recent evidence is mounting to suggest that the brain operates at-or-near criticality (Jones et al., 2023; Linkenkaer-Hansen et al., 2001; O’Byrne and Jerbi, 2022; Palva et al., 2013), a point poised between order and disorder that provides maximal flexibility, stability, and responsiveness to the environment (Cocchi et al., 2017; Hengen and Shew, 2025). This is highlighted by studies finding that brain criticality is associated with optimal cognitive functioning (Chang et al., 2026; Hengen and Shew, 2025; Müller et al., 2025; Xin et al., 2025), suggesting that critical dynamics support neural computational efficiency.

Previous research has demonstrated that critical dynamics within the brain is modulated by meditative practice, although the directionality of results are mixed. A recent study found that long range temporal correlations (LRTCs), a method to infer scale-free activity (a signature of criticality), in the theta (4-8Hz), alpha (8-13Hz), and beta (13-30Hz) bands decreased within jhāna meditation compared to mindfulness meditation (Mago et al., 2025) However, LRTCs were found to increase in the gamma (30-45Hz) band^1^. Conversely, another study found that gamma band LRTCs decrease in both focused attention (Shamatha) and open monitoring (Vipassana) practices relative to resting-state (Pascarella et al., 2025), with no significant findings in other frequency bands using cluster-based permutation tests. Further, in a separate study, LRTCs across frequency bands were found to decrease in expert practitioners, but not controls, when meditating compared to a resting-state baseline (Irrmischer et al., 2018). Interestingly, when following participant’s during one-year of meditation training, resting-state LRTCs increased, suggesting a distinct state/trait alteration as a function of meditative practice (Cahn and Polich, 2006). The differing directionality of these effects may be due to the distinct practices studied, coupled with a lack of robust phenomenological measures.

As meditation is defined as a state with significant alterations in subjective experience, it is increasingly important to apply methodologies that measure phenomenology and integrate this with neural changes within contemplative neuroscience (Lutz et al., 2024; Timmermann et al., 2023). This is particularly pertinent in phenomenologically-rich ASCs, such as meditation and psychedelics, which have temporally extended shifts in subjective experience that accompany the practice/pharmacodynamics of the substance (Abdoun et al., 2024; Jachs et al., 2022; Lewis-Healey et al., 2026; Timmermann et al., 2019). The integration of phenomenology and neural dynamics is dubbed neurophenomenology (Lutz, 2002; Lutz and Thompson, 2003; Varela, 1996), which provides a more fine-grained understanding of how neural changes are related to experiential changes, arguably one of the overarching phenomenon of study within the cognitive neuroscience of consciousness.

To investigate how critical brain dynamics change as a function of TM practice and expertise, we employed detrended fluctuation analysis (DFA) and functional excitation/inhibition (fEI) methodologies in an EEG dataset of expert TM practitioners and a control group. DFA provides an estimate of scale-free activity through the quantification of LRTCs. FEI analyses provide further information to DFA by indicating whether a system changes in a sub- or super-critical direction, with recent research indicating that the psychedelic N,N-dimethyltryptamine (DMT) induces sub-critical dynamics relative to placebo (Irrmischer et al., 2026). We utilise state-based analyses, comparing the broad state of TM to pre- and post-resting-state scans within-group, and also comparing TM in expert practitioners compared to an age and sex-matched control group. Beyond this, we applied neurophenomenological analyses, collecting time-resolved subjective effort using Temporal Experience Tracing (TET; Chandia-Jorquera et al., 2026; Jachs, 2021; Lewis-Healey et al., 2026, 2024; Niedernhuber et al., 2024), a methodology that allows participant’s to retrospectively report on dynamic experiential changes during meditation (see Methods). We related time-resolved changes in subjective effort to changes in LRTCs and fEI, to provide a more fine-grained understanding of how critical dynamics are associated with the phenomenology of TM.

## METHODS

### Participants

Data from this study has been previously published (Chandia-Jorquera et al., 2026). Briefly, participants were experienced TM meditators (N=33; 21 male; M=40.3 years old, SD=6.2) and a control group (N = 33; 21 male; M = 38,9 years old; SD = 7.5). All participants within the TM group had a minimum of four years of regular meditation practice (twice a day, 20 minutes per session; M = 12.9 years of practice; SD = 7.7). The control group (C) comprised of sex-and age-matched (±3 years) participants who had not practiced TM before. No participants had any chronic mental health, sleep-related or neurological medical problems. Meditators were recruited by the David Lynch Foundation UK, in collaboration with the Maharishi Foundation, from across the United Kingdom. Healthy controls were recruited through the University of Cambridge online recruitment system. All participants signed an informed consent, and the Cambridge Psychology Research Ethics Committee approved the experimental protocol.

### Temporal experience tracing (TET)

TET is a retrospective phenomenological methodology that preserves the dynamics of subjective experience. It has been applied in a variety of studies to investigate the phenomenological dynamics of altered states of consciousness (Chandia-Jorquera et al., 2026; Jachs, 2021; Jachs et al., 2022; Lewis-Healey et al., 2026, 2024), as well as clinical issues (Gernert et al., 2024; Niedernhuber et al., 2024), and task-based paradigms (Holgado et al., 2024, 2025). To apply the TET methodology, dimensions of subjective experience that are most relevant to the study phenomenon are identified, relating to the more fine-grained contents of consciousness (e.g., affective or attentional aspects of the experience). Following the completion of a session relating to the phenomenon of study, participants complete a retrospective trace of the intensity of each specific dimension of experience. For this study, participants were asked about the subjective effort they exerted during the meditation. Specifically, the participants were asked to trace: “How much did you have to focus/concentrate during the session?”. High values indicated a very effortful moment during the meditation/control condition, while low values indicated effortlessness.

### Experimental design and procedure

#### Transcendental Meditation

For the TM group, participants began with a 10-minute pre-meditation resting-state (pre-RS) scan, during which they were instructed to keep their eyes closed and not to engage in formal meditation. Following this, participants were instructed to engage with the practice, whereby they had to mentally repeat their mantra with their eyes closed for 30-minutes (i.e., the TM condition). Following this, participants were introduced to the process of TET, as well as the definition of the dimensions. After completing the Temporal Experience Traces, the participants completed a 10-minute post-meditation resting-state (post-RS) scan, with the same instructions as the pre-RS condition. The experimenter indicated the beginning and end of each condition.

#### Non-meditative control task

For the control group, the experimental design remained the same for the pre-RS and post-RS conditions and TET instructions. However, for the 30-minute condition, control participants were instructed to mentally count in increments of 1 with their eyes closed, at their own pace, and to restart the count at any time if they lost track. The purpose of this control task was to engage participants in cognitive processes distinct from TM but with similar automaticity and cognitive control as during mantra repetition performed by the TM group.

## TET data preprocessing

Temporal Experience Traces were digitised from scanned hand-drawn timeseries using a custom python script. The script converted the image coordinates from the TET data to graph coordinates based on the orientation of the x- and y-axes. The TET data were manually checked to ensure the vectors were similar to the hand-drawn traces. All traces were assigned to 60 equally spaced time points (one timepoint representing 30s of subjective experience), and scaled on a range of 0-1.

### EEG data preprocessing

EEG signals were recorded using a 128-channel HydroCel Geodesic Sensor Net (GSN 128 1.0) and a GES400 Electrical Geodesics amplifier, sampled at 250 Hz with NetStation software (EGI, USA). During recording and analysis, the average across electrodes was used as the reference electrode. Following data collection, continuous EEG data was band-pass filtered between 0.5–45 Hz using a zero-phase finite impulse response (FIR) filter implemented in MNE-Python. Data were subsequently visually inspected, and noisy channels were identified manually. Rejected channels 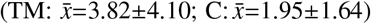 were reconstructed using spherical spline interpolation. Following bad-channel interpolation, independent component analysis (ICA) was performed using the MNE-Python implementation of FastICA. Components were manually inspected, and those judged to reflect non-neural artefacts were excluded from the data 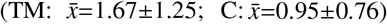 and the cleaned signal reconstructed using the remaining components. Following ICA-based artefact correction, continuous EEG recordings were segmented into consecutive non-overlapping 5-s epochs using MNE-Python. An automated artefact-screening procedure was then applied to identify epochs containing large-amplitude noise. Epochs were flagged when more than 50% of EEG channels exhibited voltages exceeding ±250 µV at any time point within the epoch. Flagged epochs were marked as bad and subsequently reviewed during visual inspection, and any epochs that were deemed as too noisy were removed from further analysis 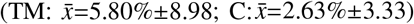. All epochs were then manually inspected, allowing additional artefactual segments to be marked for exclusion where necessary.

### Criticality metrics

#### Long range temporal correlations (LRTCs)

Detrended fluctuation analysis (DFA; Peng et al., 1994) enables the estimation of long range temporal correlations (LRTCs) - scale-free temporal dynamics within physiological timeseries. DFA was applied to the amplitude envelope of each of the canonical frequency bands (δ: 1-4Hz; θ: 4-8Hz; α: 8-13Hz; β: 13-30Hz), as well as broadband signals (BB: 1-30Hz). We applied DFA on fixed 3-minute windows on all available conditions (i.e., Pre-Rs/M/Post-RS conditions that had 36 contiguous 5s epochs). To do this, 3-minute EEG signal windows were band-pass filtered into canonical frequency bands using a zero-phase finite impulse response (FIR) filter. The amplitude envelope was then obtained using the Hilbert transform on the bandpassed signal. To compute DFA, the cumulative sum of the amplitude envelope is calculated, followed by computation of the root mean squared (RMS) fluctuation. This process is conducted for window sizes (with 50% overlap) of varying lengths. A scaling exponent is then fit via linear regression between the log-log plot of window sizes and the RMS fluctuation for that window size. Minimum window sizes were set at 5s for delta, theta, and broadband signals, 3.981s for alpha-band signals, and 2.238s for beta-band signals, while maximum window sizes were set at 30s for all frequency bands.

#### Functional excitation/inhibition ratio (fEI)

To assess whether brains exhibited sub-critical, critical, or super-critical dynamics, we applied the functional excitation/inhibition ratio (fEI) algorithm. The fEI algorithm is based on a computational model of neuronal oscillations (Poil et al., 2012); a 50}50 grid of integrate-and-fire excitatory and inhibitory neurons, which can produce signals statistically similar to human M/EEG recordings. Within this model, a balance in the E/I ratio produces critical dynamics, with DFA exponents approaching one (Bruining et al., 2020). Estimating sub-critical, critical, or supercritical dynamics from the DFA alone within real human physiological data is not possible, as scaling exponents are organised as an inverse u-shape around the E/I ratio. To circumvent this, the fEI algorithm was developed, demonstrating that distinct associations between the RMS fluctuation function of smaller window sizes and oscillatory amplitudes yielded an estimate of sub-critical (positive association), critical (no significant association), or super-critical (negative association) dynamics (see Figure 1 for a schematic illustration). Thus, the fEI algorithm provides a finer-grained estimate of how the brain’s critical dynamics changes as a function of meditation or meditative expertise.

**Figure 1:**
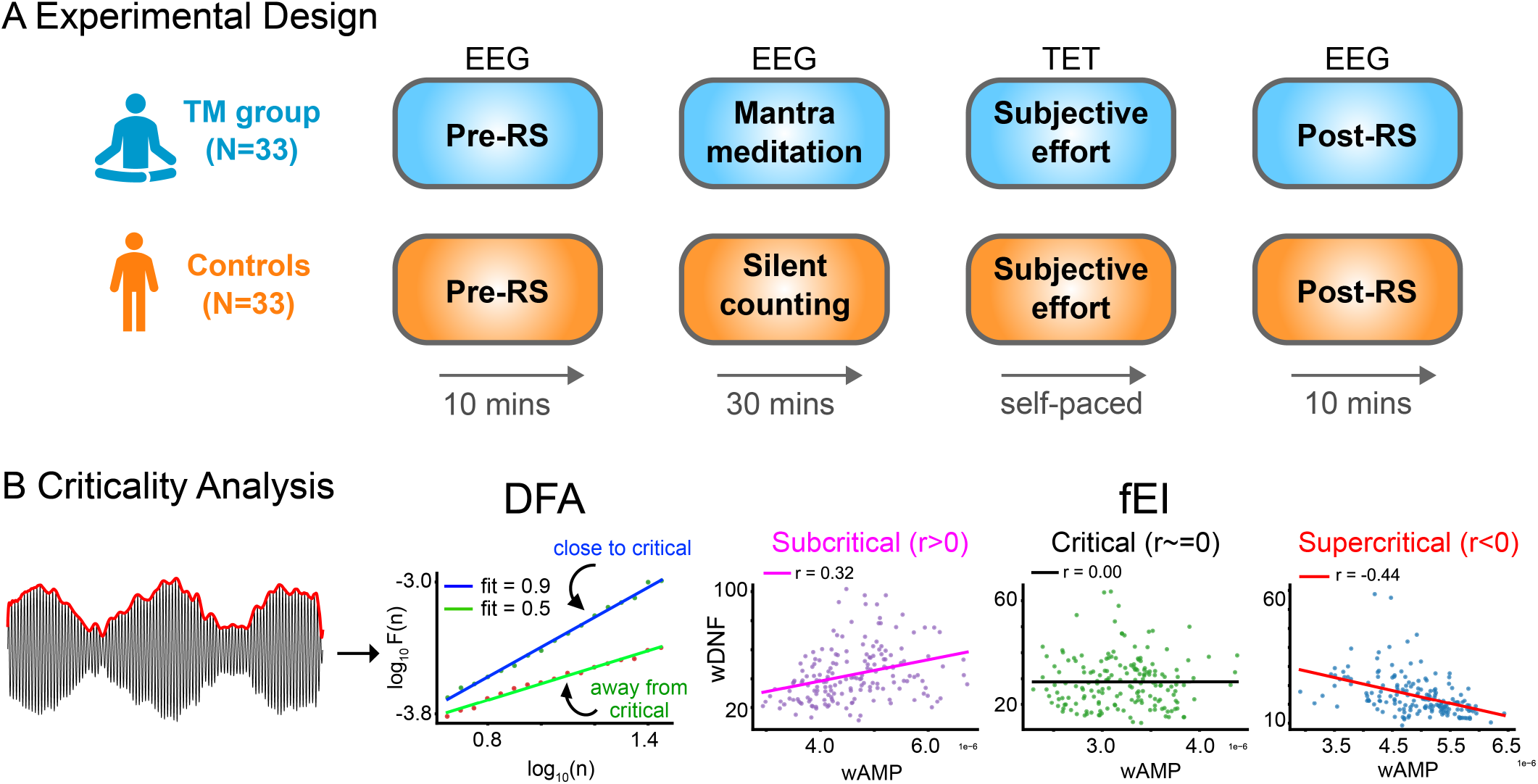
Experimental design and criticality analyses. **(A)** Participants were either experienced TM practitioners (N=33) or meditation-naive control participants (N=33). TM practitioners underwent a Pre-meditation resting-state (Pre-RS) EEG scan, followed by a 30-minute TM session. After the TM session, the participant reported on their subjective effort over time, using Temporal Experience Tracing (TET). Succeeding this, participants underwent a post-meditation RS scan (Post-RS). For control participants, the experimental design was the same, but instead of TM, the controls counted silently in their head during the 30-minute session. **(B)** From the amplitude envelope of the bandpass filtered signal (in red, left), detrended fluctuation analysis (DFA) and the functional excitaion/inhibition ratio (fEI) was computed. DFA plots the association between the window sizes and the detrended normalised fluctuation function on a log-log scale, with values close to 1 being close-to-critical. FEI plots the association between smaller windowed amplitudes (wAMP) and the windowed detrended normalised fluctuation function (wDNF). Positive linear associations indicate subcritical dynamics, no discernible associations indicate critical dynamics, whereas negative linear associations indicate supercritical dynamics.

FEI was estimated following the procedure from Bruining et al. (2020). Briefly, the signal was bandpass filtered into canonical frequency bands, and the amplitude envelope of the signal was extracted using the Hilbert Transform (same procedure as DFA). Following this, the cumulative sum of the amplitude envelope is calculated, segmented into 80% overlapping 5s windows, normalised by dividing each value by the mean of the original window, and detrended. The RMS function is then computed on each detrended, normalised window, and the Pearson’s correlation between the oscillatory amplitude and the windowed RMS fluctuation is computed and subtracted from 1, providing the fEI value. Values close to 1 (i.e., no association between oscillatory amplitudes and detrended normalised fluctuations) indicate critical dynamics, whereas negative values indicate subcritical dynamics, and positive values indicate supercritical dynamics. To reduce the influence of extreme windows, outliers were identified separately for the windowed amplitude and windowed detrended normalised fluctuation using the generalized extreme studentized deviate (GESD) procedure (Rosner, 1983) with a significance level of 0.05. A maximum of 2.5% of windows per measure (approximately 5% combined) were eligible for removal. Outlier windows identified were excluded before computing the fEI estimate. Following Bruining et al. (2020), fEI values were considered valid only for channels with DFA exponents greater than 0.6.

#### TET-criticality metric alignment

To conduct neurophenomenological statistical analyses, time-resolved subjective effort (collected via Temporal Experience Traces) were aligned with corresponding DFA and fEI values. To do this, for each EEG window that criticality was computed on (i.e., 3-minutes), TET epochs whose temporal centre fell within the EEG window were identified and assigned to that window. The phenomenological value associated with each criticality window was then calculated as the mean of the corresponding TET epochs for subjective effort. Thus, a single value of subjective effort was assigned to each corresponding criticality metric.

#### Statistical Analyses

To investigate spatially distributed differences in criticality metrics, cluster-based permutation tests were performed on channel-wise DFA and fEI values. For within-group comparisons between conditions (Pre-RS, Meditation, and Post-RS), paired posthoc cluster-based permutation tests were conducted if a repeated-measures ANOVA was found significant on the whole-brain average values for the corresponding criticality metric. For between-group comparisons, independent-samples cluster-based permutation tests were conducted. Clusters were formed from neighbouring electrodes exceeding a predefined cluster-forming threshold (two-tailed (*p*<0.01) for between-group analyses; an adaptive threshold was used for within-group one-sample tests). Statistical significance was determined using 5,000 random permutations, with cluster-level p-values calculated relative to the permutation-derived null distribution of maximum cluster statistics. Cluster-level p-values were corrected for multiple comparisons across frequency bands using the Benjamini–Hochberg false discovery rate (FDR) procedure (Benjamini and Hochberg, 1995).

For the within-meditator criticality cluster-based permutation tests, a subset of the participants were excluded from the analysis as at least 3-minutes of contiguous EEG data (i.e., no epoch removal) had to be included for each of the conditions for the reliable computation of the criticality metrics. Therefore, 21 participants were included for all DFA metrics, while, for the fEI analyses, 18 were included for the beta band, 19 for the theta and alpha bands, and 20 for the delta band and broadband. The fEI analyses were more variable in their participant inclusion as fEI was not computed on channels with DFA exponents greater than 0.6, as fEI is only a valid measure when there are underlying critical dynamics (Bruining et al., 2020). We therefore used a threshold of at least 50% of the channels had to have non-NaN fEI values to be included in the analysis.

We employed linear mixed models to investigate the difference between groups with regards to the intensity of subjective effort during the meditation/control condition, whilst providing subject as a random effect. To investigate the association between criticality metrics and subjective effort, linear mixed models were also utilised. To compare the results of distinct criticality-effort associations (e.g., comparing α-LRTCs to β-LRTCs), all variables were standardised to have mean of 0 and variance of 1, thus allowing the coefficients to provide a comparable indication of the strength of association, similar to effect sizes (Lorah, 2018). Prior to the computation of any linear mixed models, all criticality metrics that were above or below 2.5SDs of the mean were removed from statistical analysis.

#### Hypotheses

Based on a recent theoretical framework of the consequences of meditation on neural activity (Laukkonen, 2026), we hypothesised that TM practitioners would have increased LRTCs and closer-to-critical regimes during TM relative to pre- and post-RS. Further, we also hypothesised that TM practitioners would have higher LRTCs and fEI values closer to criticality (i.e., nearer to 1) than controls when practicing TM compared to the control counting condition. Further, we hypothesised that effort would be significantly associated with critical brain dynamics.

#### Software and analyses

Statistical and signal processing analyses were all applied in Python (3.11.8). Criticality was computed using the Critical Oscillations toolbox (https://github.com/Critical-Brain-Dynamics/crosci/).

## RESULTS

### Transcendental Meditation is associated with increased LRTCS within-meditators

To investigate whether there was a significant difference in long range temporal correlations (LRTCs) and the functional excitation/inhibition ratio (fEI) within meditation compared to resting-state, we compared LRTCs and fEI in the canonical frequency bands (δ: 1-4Hz; θ: 4-8Hz; α: 8-13Hz; β: 13-30Hz), and broadband signals (BB: 1-30Hz), between the three conditions (pre-RS, M, post-RS). To compare the differences between conditions, we first ran a repeated-measures ANOVA on each whole-brain metric. If there was a significant difference between a metric (after FDR correction), we applied posthoc pairwise cluster-based permutation paired t-tests between each condition.

Within the TM group, α-LRTCs (*F*(2,40)=6.22, *p*=0.011, 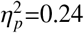) and β-LRTCs (*F*(2,40)=6.79, *p*=0.011,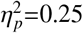) were significantly different between all three conditions, while δ-LRTCs (*F*(2,40)=0.59, *p*=0.56, 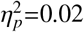, θ-LRTCs (*F*(2,40)=2.71, *p*=0.13,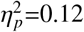), and BB-LRTCs (*F*(2,40)=0.94, *p*=0.40,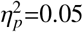) were not significantly different between conditions. Posthoc cluster-based permutation tests revealed a significant increase in α-LRTCS during TM relative to pre-RS, comprising a large positive cluster spanning 104 channels barring centro-posterior sites (*p*=0.019). A significant cluster was observed when comparing TM to post-RS, spanning 94 channels barring centro-posterior sites (*p*=0.035; Figure 2A). No significant differences were observed between pre-RS and post-RS. Posthoc cluster-based permutation tests revealed heightened β-LRTCs within TM compared to both pre- and post-RS. TM vs. pre-RS contrasts yielded a large cluster involving 116 channels (*p*=0.019), while the TM vs. post-RS comparison yielded a significant cluster involving 72 fronto-central and occipital channels (*p*=0.035). No additional β-LRTC clusters survived FDR correction, and no significant differences were observed between pre-RS and post-RS.

**Figure 2:**
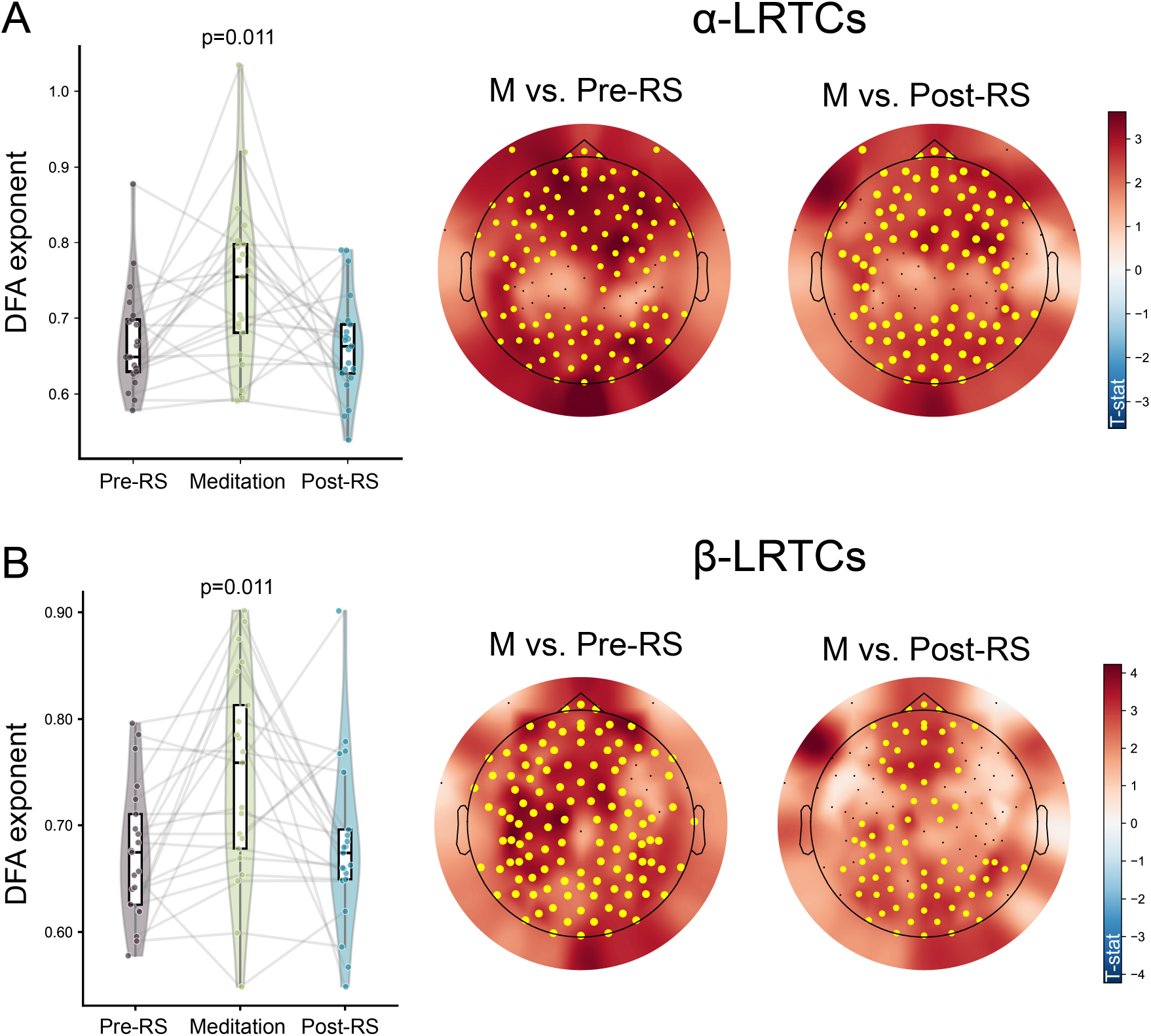
Transcendental Meditation is associated with an increase in α-LRTCs and β-LRTCs compared to pre- and post-resting state.. **(A)** (Left) Whole-brain averages of α-LRTCs (8-13Hz) for the TM group across each condition. Lines connect whole-brain average values for the same participant across all three conditions. A repeated-measures ANOVA revealed a significant difference between conditions. (Right) Posthoc cluster-based permutation tests indicated that α-LRTCs were significantly higher during TM compared to both pre- and post-RS, indicating closer-to-critical temporal oscillatory dynamics in 8-13Hz. **(B)** (Left) As above, whole-brain averages of β-LRTCs (13-30Hz) for the TM group across each condition. Lines connect whole-brain average values for the same participant across all three conditions. A repeated-measures ANOVA revealed a significant difference between conditions. (Right) Posthoc cluster-based permutation tests indicated that β-LRTCs were significantly higher during TM compared to both pre- and post-RS, indicating closer-to-critical temporal oscillatory dynamics in 13-30Hz.

Repeated-measures ANOVA revealed that no fEI metrics were significantly different between the three conditions for δ-fEI (*F*(2,40)=1.08, *p*=0.43,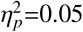), β-fEI (*F*(2,36)=0.69, *p*=0.51,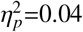), and BB-fEI (*F*(2,40)=2.51, *p*=0.16,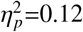). θ-fEI (*p*=0.03) and α-fEI (*p*=0.02) were significantly different between conditions, however, neither θ-fEI (*F*(2,38)=3.97, *p*=0.07,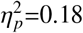) nor α-fEI (*F*(2,38)=4.30, *p*=0.07,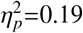) survived FDR correction (see Supplementary Figure S1 for whole-brain fEI comparisons between conditions). Therefore, no posthoc cluster-based permutation tests were conducted.

### No state-based difference in critical brain dynamics between meditators and controls

To test whether there were significant differences between the TM and control groups, we ran cluster-based permutation independent-samples t-tests between the TM and counting conditions in all DFA and fEI metrics, with each metric baseline-corrected relative to pre-RS values. However, contrary to our predictions, no significant clusters were found in any frequency band for both DFA and fEI metrics (see Supplementary Figures S2 and S3).

### Subjective effortlessness is robustly associated with critical brain dynamics in meditators, but not controls

Due to previous findings highlighting that subjective effort is associated with critical brain dynamics (Fagerholm et al., 2015; Kardan et al., 2020; Lim et al., 2026), and that effortlessness is a major component and phenomenological goal of TM (Farrow and Hebert, 1982; Mahone et al., 2018; Tang et al., 2022) and meditative practices more broadly (Garrison et al., 2013; Jachs, 2021; van Lutterveld et al., 2017), we investigated the association between time-resolved subjective effort and criticality metrics.

First, we found that the control group reported significantly higher subjective effort^2^ across the session compared to the TM group (β=0.82, *SE*=0.178, *z*=4.62, *p*<0.001, 95% CI [0.473,1.170]), indicating that meditators experienced more effortless states during TM compared to controls conducting the counting condition (Figure 3A). Building on this, to quantify the temporal divergence in effort shown in Figure 3A, we fit a linear mixed model with time, group, and their interaction as fixed effects and participant as a random intercept. Results indicated that effort significantly decreased over time for the TM group (β=-0.009, *SE*=0.001 *z*=-15.42, *p*<0.001), and the interaction effect indicated that controls significantly differed in this change in effort over time (β=0.020, *SE*=0.001 *z*=23.65, *p*<0.001), indicating that effort increased over time in controls relative to meditators.

**Figure 3:**
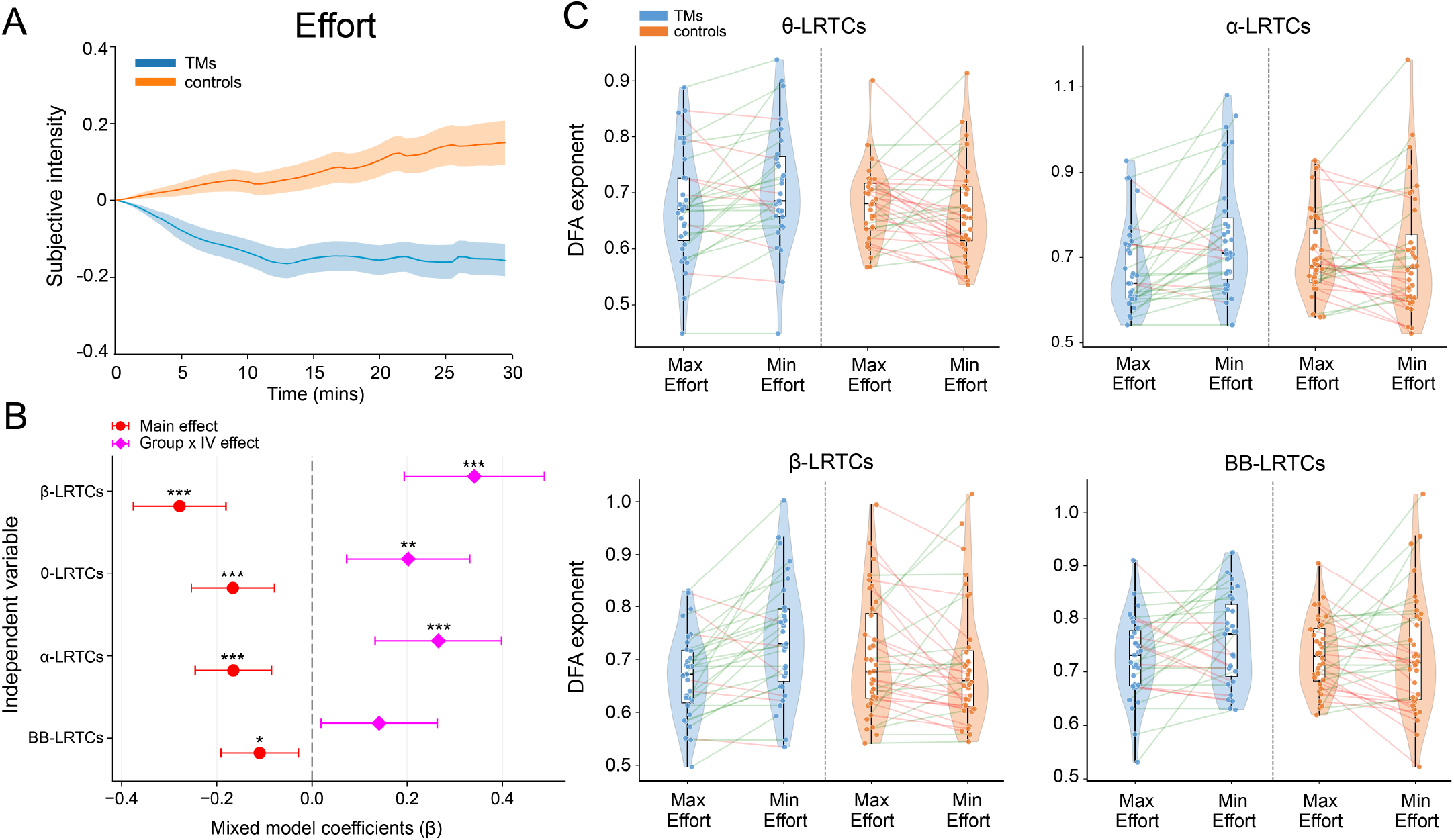
Subjective effortlessness in meditation is robustly associated with higher LRTCs within meditators, but not control participants.. **(A)** Time-resolved subjective effort for meditators (blue) and controls (orange) during the meditation/counting condition. Dark lines indicate the mean value over all participants, while shaded areas indicate the standard error of the mean. Each participant’s values are baseline-corrected relative to the first datapoint of the session. **(B)** Main (red) and group interaction (purple) effects for linear mixed models investigating the association between criticality metrics and subjective effort. Error bars represent the 95% confidence intervals for the corresponding coefficients. The negative main effects indicate an increase in LRTCs are associated with a decrease in subjective effort. Positive interaction effects indicate that the association between subjective effort and LRTCs is significantly weaker for control participants. **(C)** Subject-level differences in LRTCs across significant frequency bands for the corresponding minimum and maximum subjective effort of each session for meditators (in blue) and controls (in orange). Green lines indicate higher LRTCs when a participant is in a comparatively effortless state. \*\*\**p*<0.001; \*\**p*<0.01; \**p*<0.05

Following this, we used linear mixed models to investigate whether there was a significant association between subjective effort and criticality metrics, and if there was a group-level interaction effect. Baseline-corrected effort was significantly associated with LRTCs across multiple frequency bands. Effort showed an inverse association with θ-LRTCs (β=-0.17, *SE*=0.044, *z*=-3.73, *p*<0.001, 95% CI [−0.253, −0.079]), α-LRTCs (β=-0.17), *SE*=0.041, *z*=-4.04, *p*<0.001, 95% CI [−0.245, −0.085]), β-LRTCs (β=-0.28, *SE*=0.050, *z*=-5.60, *p*<0.001, 95% CI [−0.375, −0.181]), and BB-LRTCs (β=-0.11, *SE*=0.041, *z*=-2.65, *p*=0.02, 95% CI [−0.191, −0.029]). Thus, lower subjective effort was associated with increased LRTCs, and hence closer-to-critical brain dynamics, across these frequency ranges. No significant main (or group-level interaction) effects were found across any fEI metrics (see Supplementary Table S1 for full results).

Significant group interactions were found for *θ*-LRTCs, *α*-LRTCs, and *β*-LRTCs, indicating that the relationship between effort and LRTCs differed between meditators and controls. Specifically, the inverse association between subjective effort and *θ*-LRTCs was significantly attenuated in controls relative to meditators (*β*=0.20, *SE*=0.066, *z*=3.07, *p*=0.007, 95% CI [0.073, 0.331]). Similar interaction effects were observed for *α*-LRTCs (*β*=0.27, *SE*=0.068, *z*=3.92, *p*<0.001, 95% CI [0.133, 0.398]) and *β*-LRTCs (*β*=0.34, *SE*=0.075, *z*=4.54, *p*<0.001, 95% CI [0.194, 0.488]). The significant group interaction in BB-LRTCs, however, did not survive FDR correction (*β*=0.14, *SE*=0.06, *z*=2.27, *p*=0.058, 95% CI [0.019, 0.263]). These positive and significant interaction coefficients indicate that the inverse association between effort and LRTCs was substantially stronger in the TM group than in controls. In other words, effortlessness was associated with increased LRTCs in meditators, whereas this relationship was markedly weaker (and in some bands absent) in controls (Figure 3B&C).

### Effort-criticality associations are not explained away by time spent in meditation

As time-resolved effort diverges between meditators and controls - decreasing/increasing respectively over time (relative to baseline; Figure 3A) - we also investigated whether there was a significant association between the time spent during meditation/counting and critical brain dynamics. When running linear mixed models evaluating the association between time spent in meditation and criticality metrics, only *α*-LRTCs (*β*=0.25, *SE*=0.054, *z*=4.63, *p*<0.001, 95% CI [0.144, 0.357]) and *β*-LRTCs (*β*=0.23, *SE*=0.045, *z*=5.16, *p*<0.001, 95% CI [0.144, 0.321]) showed positive significant associations with time spent in counting/meditation. However, neither group-level interaction effects were significant. It therefore appeared that both *α*- and *β*-LRTCs increased as a function of time within counting/meditation across both groups.

To explore whether time spent in meditation could explain away the effort-criticality associations within both *α*- and *β*-LRTCs (i.e., whether the effort-criticality associations remained), we ran two separate linear mixed models with both subjective effort and time spent in meditation as independent variables, and the corresponding DFA metrics as the dependent variable, with an effort x group interaction, and subject as a random effect. We found that effort remained a strong predictor of *α*-LRTCs (*β*=-0.47, *SE*=0.09, *z*=-4.97, *p*<0.001, 95% CI [−0.657, −0.285]), as did the effort x group interaction term (*β*=0.43, *SE*=0.13, *z*=3.45, *p*<0.001, 95% CI [0.187, 0.679]), and time (*β*=0.14, *SE*=0.03, *z*=4.29, *p*<0.001, 95% CI [0.077, 0.204]). Similarly, effort remained a strong predictor of *β*-LRTCs (*β*=-0.50, *SE*=0.09, *z*=-5.71, *p*<0.001, 95% CI [−0.672, −0.328]), alongside a significant effort × group interaction (*β*=0.45, *SE*=0.12, *z*=3.89, *p*<0.001, 95% CI [0.223, 0.675]) and a significant positive effect of time (*β*=0.14, *SE*=0.03, *z*=4.92, *p*<0.001, 95% CI [0.087, 0.202]). These models had a lower Bayesian Information Criterion (BIC) when compared to simply modelling the association between time spent in meditation/counting and *α*-LRTCs (ΔBIC=6.85) and *β*-LRTCs (ΔBIC=15), providing strong evidence that including an effort x group fixed effect interaction fits the data better than simply including time as a fixed effect (Shen and González, 2021). These findings suggest that the inverse association between subjective effort and LRTCs persisted even after accounting for temporal changes across the meditation/counting session, and that this relationship remained substantially stronger in meditators than in controls. Thus, subjective effort within TM is robustly associated with critical dynamics across the frequency spectrum that can not be explained simply through time spent in meditation, highlighting the strength of our neurophenomenological approach.

### Group-level differences in beta-band LRTCs emerge when applying phenomenologically-guided analysis

Building on the fact that there were robust associations between subjective effort and LRTCs across frequency bands, with significant differences between meditators and controls, we finally explored whether there were channel-level differences between groups using cluster-based permutation tests. However, rather than a purely state-based analysis as conducted earlier, we compared criticality metrics in the lowest available values of subjective effort^3^ for each participant’s meditation/counting condition. A significant fronto-central cluster for *β*-LRTCs, formed of 20 channels, survived FDR correction (*p*=0.042), where baseline-corrected (M - pre-RS) LRTCs were higher in meditators compared to controls (Figure 4). No other clusters across any DFA and fEI metrics survived FDR correction. As the lowest-effort windows selected for this analysis may occur at different times in the session between groups, we conducted a follow-up analysis to determine whether this temporal difference could account for the observed *β*-LRTCs effect using averaged whole-brain values. An Ordinary Least Squares (OLS) linear regression model predicting the onset time of the lowest-effort window from group indicated that controls were at their lowest-effort period significantly earlier than meditators (*β*=-376.09s, *t*=-2.79, *p*=0.007). We therefore fitted a separate OLS linear regression model for *β*-LRTCs, with whole-brain *β*-LRTCs predicted by window onset time, group, and their interaction. Neither the main effect of window onset time nor the time × group interaction reached significance for *β*-LRTCs (time: *p*=0.416; time × group: *p*=0.274). These findings suggest that, although the lowest-effort windows occurred at different times between groups, the temporal position of the selected windows did not explain the observed group differences in *β*-LRTCs. In sum, we found between-group differences within meditation when guiding the analysis based on subjective effort, while group-level differences did not emerge when using a state-based approach that ignores phenomenology. This further highlights the strength of our employed neurophenomenological approach.

**Figure 4:**
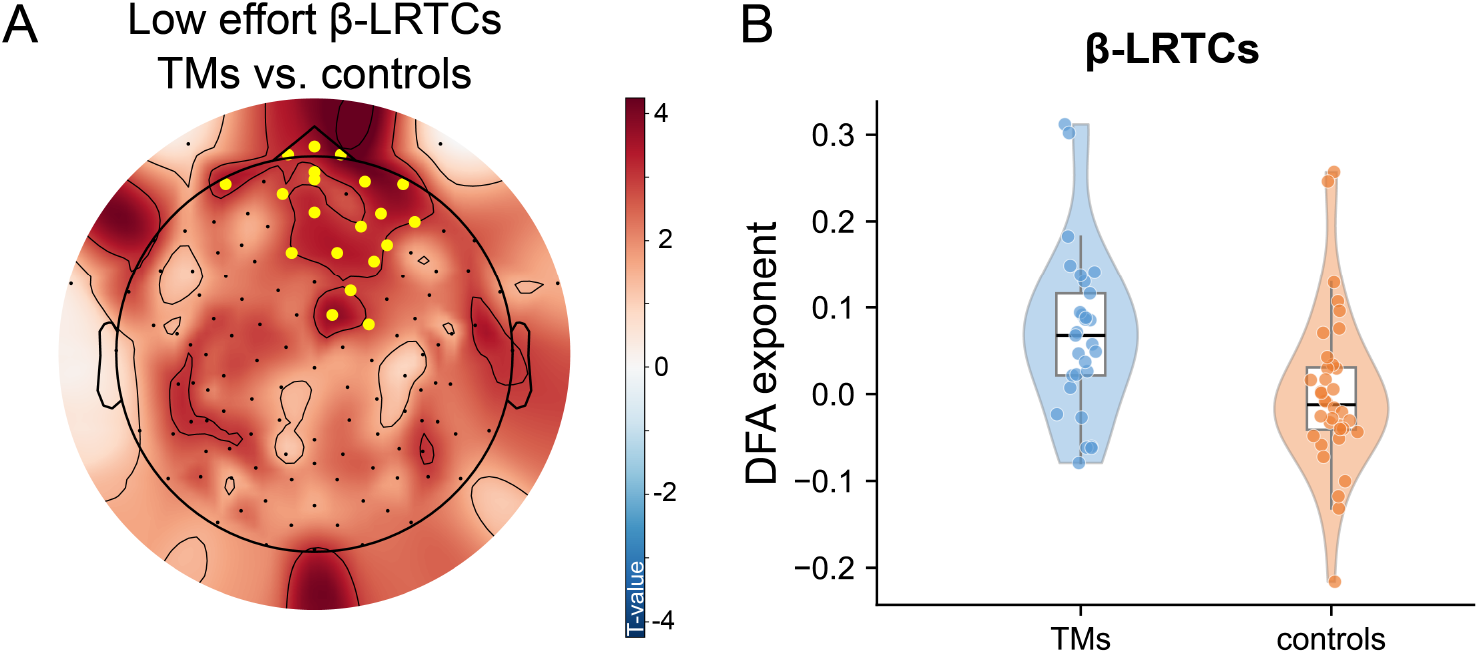
Neurophenomenological analyses revealed that meditators have significantly higher fronto-central *β*-LRTCs than controls during subjectively low-effort moments of Transcendental Meditation, highlighting group-level differences in critical brain dynamics. **(A)** Cluster-based permutation tests revealed that *β*-LRTCs were significantly higher in the TM group for fronto-central electrodes (*p*=0.042) during subjectively low-effort periods within the meditation/counting conditions. Criticality metrics were baseline-corrected relative to the within-participant pre-RS values. **(B)** Whole-brain average *β*-LRTCs during subjectively low effort periods within the meditation/counting condition. As the LRTCs are baseline-corrected, a value of 0 indicates no change between pre-RS and the subjectively low effort meditation/counting condition.

## DISCUSSION

Subjective effortlessness is a hallmark phenomenological goal of a variety of meditative practices, which may, in turn, be related to an unconstrained and flexible mode of cognition and neural dynamics. To test this, we investigated changes in critical brain dynamics of expert TM practitioners and control participants using detrended fluctuation analysis (DFA) and the functional excitation/inhibition ratio (fEI) with high-density EEG. While fEI values across frequency bands were not significantly different within-meditators across conditions, or between-groups, we found that *α*-, and *β*-LRTCs in the meditative state were significantly higher compared to pre- and post-RS scans within TM practitioners. However, using this state-based analysis, we found no significant differences in LRTCs between TM practitioners and controls. Applying a neurophenomenological analytical approach, we found that TM was associated with a more subjectively effortless state in comparison to the control counting condition, and that periods of effortlessness were robustly associated with increased LRTCs across a wide range of frequencies. Interestingly, this association - between LRTCs and subjective effort - was not found in control participants, as evidenced by significant group interaction effects. Finally, parsing criticality metrics based on low subjective effort in between-group cluster-based permutation tests indicated that *β*-LRTCs were significantly higher for the TM group in a fronto-central cluster compared to controls, highlighting that near-critical brain dynamics are more closely tied to subjective experience, rather than to the global state of meditation. Our results suggest that subjective effortlessness within TM is closely tied to a more flexible mode of cognition, as indexed by higher LRTCs across the broad frequency spectrum, which may ultimately support more optimal information processing and a computationally efficient neural state (Laukkonen, 2026).

Our within-group findings - that TM is associated with increases in LRTCs compared to resting-state - contrasts with some previous findings within contemplative neuroscience. For example, Irrmischer et al. (2018) found that LRTCs across frequency bands are actively suppressed during focused attention (FA) meditation, and Mago et al. (2025) found a reduction in *θ*-, *α*-, and *β*-LRTCs in jhāna meditation compared to mindfulness. We may explain these disparate results due to the distinct practices entailed within the studies. TM is unique in that it is a mantra-based practice, and does not use a sensory anchor (e.g., the breath) to induce meditative phenomenology like FA (Laukkonen and Slagter, 2021; Lutz et al., 2008) and jhāna (Brahinsky et al., 2024) meditation. This sustained attentional focus is in contrast with TM, which, if conducted correctly, involves an “effortless” repetition of the mantra, emphasising a lack of cognitive control. Indeed, our neurophenomenological results cohere with this, finding that effortless states within TM are associated with scale-free dynamics closer to a critical point. To bridge the gap between these potentially distinct results, we strongly suggest the use of methodologies to identify and link changes in both neural *and* phenomenological dynamics across meditative disciplines.

Beyond the quantification of LRTCs in EEG, a recent study used fMRI whole-brain modelling to quantify the bifurcation parameter in distinct brain networks, an operationalisation of critical dynamics (Vohryzek et al., 2025). It was found that there was a non-linear shift across resting-state networks in bifurcation parameters becoming closer to the critical point, most notably when entering the advanced ‘formless’ jhānas. The authors highlight that progression through the jhānas is phenomenologically described as a progressive “letting-go” of increasingly basic experiences (Burbea, 2025; Prest, 2026), suggesting and noting an effortlessness with this practice that accompanies expertise. Again, our results fit this narrative, showing a strong and robust relationship between subjective effortlessness and LRTCs. We therefore wish to once more emphasise the utility of neurophenomenology across the study of meditative disciplines, as the neural dynamical changes we observe here are tied to subjective changes as a function of meditation. We propose that scale-free dynamics might be a candidate neural marker of “letting-go” within meditation more broadly, rather than a coarse-grained neural marker of the meditative state itself.

This notion - that criticality may be a candidate neurophenomenological marker of meditative effortlessness - is also supported by research outside of contemplative neuroscience; a variety of studies have found modulations in critical brain dynamics in the face of cognitive effort and task complexity. For example, effortful engagement in working memory tasks have shown strong associations with deviation from criticality compared to rest (Kardan et al., 2020; Lim et al., 2026), suggesting that effortful cognition requires more computational stability and less flexibility. This is further supported by other evidence highlighting that focused attention induced by conducting a task is associated with subcritical dynamics (Fagerholm et al., 2015). While there may be spatial nuances in task-related deviations from criticality (Habibollahi et al., 2026; Lim et al., 2026), these findings are coherent with our results; transient periods in subjective effort during TM could entail an engagement in attention to focus on the mantra, as the mind may have wandered. This stability may lead to less neural computational flexibility, and therefore deviations from criticality, quantified by a reduction in LRTCs. Conversely, being on-task with the meditation - wielding an effortless and open awareness - may push the brain closer to a critical point. Future research may seek to investigate how criticality relates to other non-pharmacological ASCs, such as flow-states, characterised by an effortless attentiveness to the task at hand due to an optimal balance between skill-level and task-complexity (Alameda et al., 2022; Durcan et al., 2024; Gold and Ciorciari, 2020). Relating effortlessness to criticality metrics could serve as a neurophenomenological bridge between meditation and flow as distinct yet overlapping ASCs.

Across all neural metrics, *β*-LRTCs were most robustly associated with (i) state-based changes in TM, (ii) subjective effort, and (iii) between-group changes in low effort states during TM/counting. Recent research has found a strong association between self-dissolution (i.e., reductions in the felt sense of an embodied self) induced through meditation and reductions in oscillatory beta power (Dor-Ziderman et al., 2016; Trautwein et al., 2024). However, there is a nonlinear relationship between oscillatory power and LRTCs (Bruining et al., 2020), which indicate that this attenuation in oscillatory power due to meditative self-dissolution may not hold a straightforward relationship with the temporal structure of beta oscillations. Future research may therefore investigate whether meditation-induced modulations in selfhood influence critical dynamics. Interestingly, it may be that dynamical changes in criticality may be temporally-resolved, with scale-invariant dynamics being suppressed under self-dissolution conditions, and getting closer-to-criticality following this induction, as the dissolution of self may engender more flexible cognitive processing that may have transdiagnostic implications for a reduction of suffering (Cooke, 2026; Laukkonen, 2026; Laukkonen and Slagter, 2021).

One overarching finding within this study is that the integration of phenomenological information with neural dynamics led to stronger explanatory power of between-group differences. State-based analyses (i.e., contrasting TM versus counting), on the other hand, yielded no significant difference between groups. This emphasises the necessity of capturing (and explaining/predicting) subjective experience when studying ASCs (Lutz et al., 2024; Timmermann et al., 2023), which can demonstrably lead to a finer-grained understanding of neural dynamics and how they change as a function of ASC modulation (Jachs, 2021; Jachs et al., 2022; Lewis-Healey et al., 2026, 2024; Lutz et al., 2002; Lutz and Thompson, 2003; Varela, 1996). Further, utilising TET as the phenomenological methodology, rather than retrospective Likert Scale questionnaires, provided a more temporally-resolved pinpointing of subjective effortlessness, which is highly suitable for meditation as a temporally diffuse ASC driven by subjective fluctuations in attention and effort. We therefore encourage future researchers within the human cognitive neurosciences more broadly to work from the premise of neurophenomenology, and integrate more sophisticated phenomenological methodologies within novel studies.

Finally, we highlight some limitations and suggestions for future research. First, while TET provides a time-resolved marker of subjective effort, neurophenomenological analyses entailed the coarse-graining of the TET data, as both DFA and fEI were computed with 3-minute contiguous stretches of EEG (and TET datapoints were originally associated with 30s of data). Despite robust neurophenomenological findings, future analysis could seek to incorporate more time-resolved measures of criticality (and phenomenology) that have been developed (Lim et al., 2026; Sooter et al., 2025). Further, as EEG provides poor spatial granularity and source-localisation, future research may seek to utilise MEG, which could identify specific regions of the brain that are modulated through meditative practice, as well as reliably investigating higher frequency signals (Muthukumaraswamy, 2013), and could be used to build more advanced computational models of brain activity during meditation (Cofré et al., 2020; Patow et al., 2024).

In sum, we demonstrate that near critical brain dynamics are modulated through the practice of TM, whereby the brain operates closer-to-criticality during TM compared to resting-state. Within TM, increased LRTCs across a broad frequency range are predictive of subjective effortlessness, suggesting that the temporal structure of oscillatory activity may be a candidate neural marker for “letting-go” across a range of meditative practices. Our results ultimately contribute to a more nuanced picture of TM, and meditative practices more broadly, moving away from the notion of a meditative “state”, and towards neurally understanding the more fine-grained fluctuations in conscious experience that result.

## Data Availability

Data from this study are available from ACJ upon reasonable request, in accordance with the Department of Psychology, University of Cambridge’s data sharing policies.

## Authorship Contribution

Conceptualization and methodology: ELH, ACJ, RL; Investigation and data collection: ACJ; Data analysis: ELH; Visualization: ELH and ACJ; Writing: ELH; Reviewing and editing: ELH, ACJ, RL; Supervision: ACJ, RL; Funding acquisition: ACJ, RL.

## Funding

Collection of this data was funded by The David Lynch Foundation UK through private donors provided to A.C-J. E.L-H and R.L. are supported by a philanthropic grant from the Cillo family provided to R.L. A.C-J. is supported by a Swedish Research Council project grant (VR; 2025-03245), a Research Council of Finland project grant (RCF; 375200), an ANID/FONDECYT Regular (1240899) and ANID/FONDECYT Regular (1251273) research grants.

## Supplementary Results

### Wakefulness does not account for the effort–LRTC relationship

To determine whether the relationship between effort and LRTCs is not simply explained by fluctuations in arousal, we fit linear mixed models predicting each DFA metric from subjective effort, subjective wakefulness (also collected via TET), group, and their interactions (effort × wakefulness × group; random intercept for subject). Across all DFA metrics, wakefulness showed no robust main effect or wakefulness × group interaction, indicating that fluctuations in subjective wakefulness had no significant association with LRTCs. Further, the three-way interaction did not survive FDR correction for any frequency band (all FDR-corrected *p*>0.28). In contrast, the main effect of effort remained significant across θ-, α-, β- and BB-LRTCs, as did the effort × group interaction for θ-, α-, and β-LRTCs (all FDR-corrected *p*<0.01), with the BB-LRTCs effort × group effect showing the same direction but not surviving FDR correction (*p*=0.08). Together, these models provide little evidence that wakefulness explains the effort–LRTC association, which remains strongest for the effort and effort × group effects.

### No effect of preprocessing on LRTCs

To test whether the difference in the preprocessing metrics between groups had an influence on the DFA metrics, we fitted linear mixed models with number of interpolated channels, ICA components removed, and rejected epochs as predictors (random intercept for subject|condition). After FDR correction, none of the preprocessing variables showed a reliable association with DFA metrics across frequency bands (all *p*>0.11). This suggests that the reported group differences in LRTCs are unlikely to be driven by differences in preprocessing.

**Figure S1:**
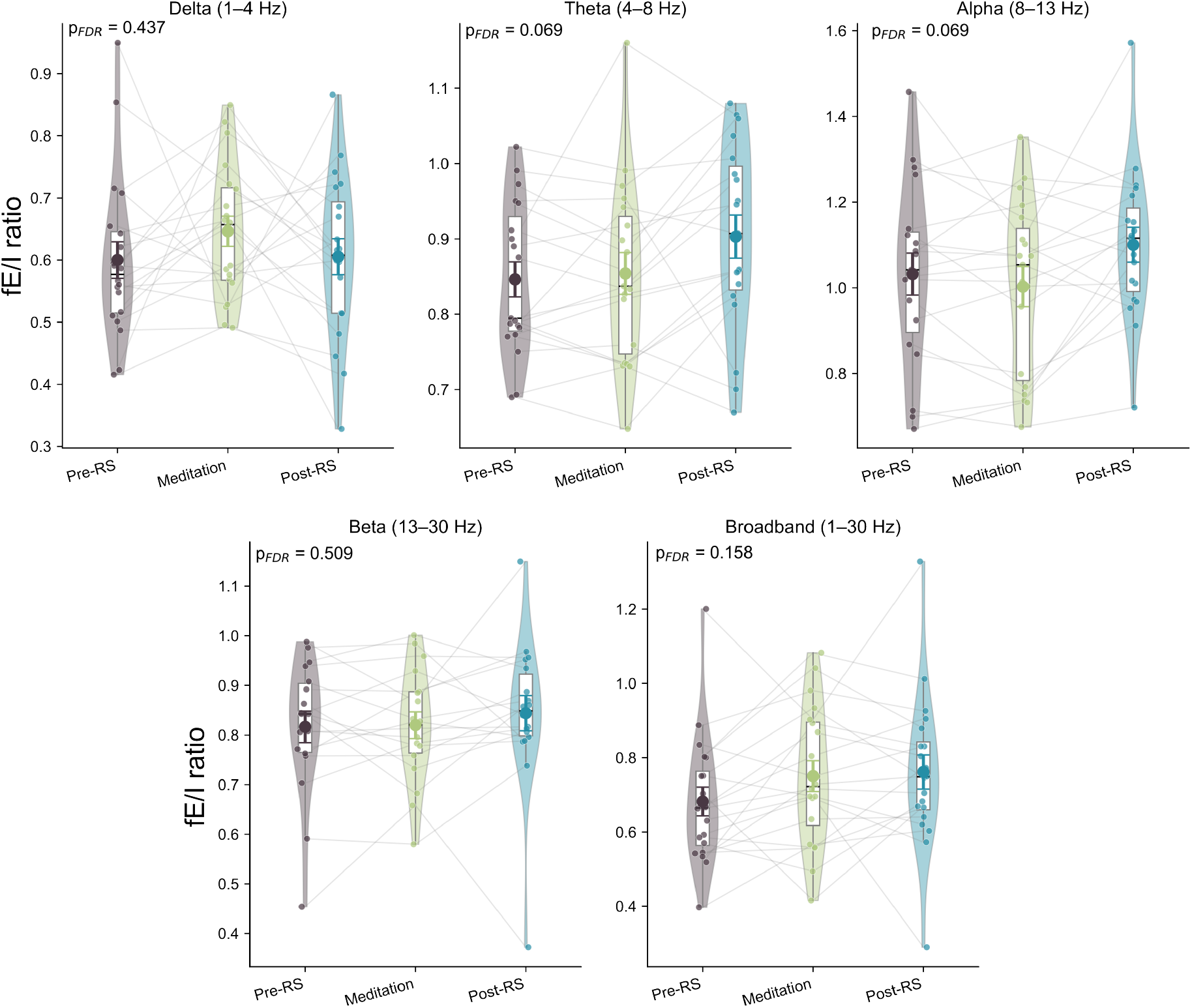
Functional Excitation/Inhibition ratio (fEI) differences between conditions within-meditators. Distribution of whole-brain values of fEI across frequency bands within-meditators. Lines between values connect the same participant across conditions. P-values (FDR corrected) of the repeated-measures ANOVA are reported in the top left of each plot for the corresponding frequency band.

**Figure S2:**
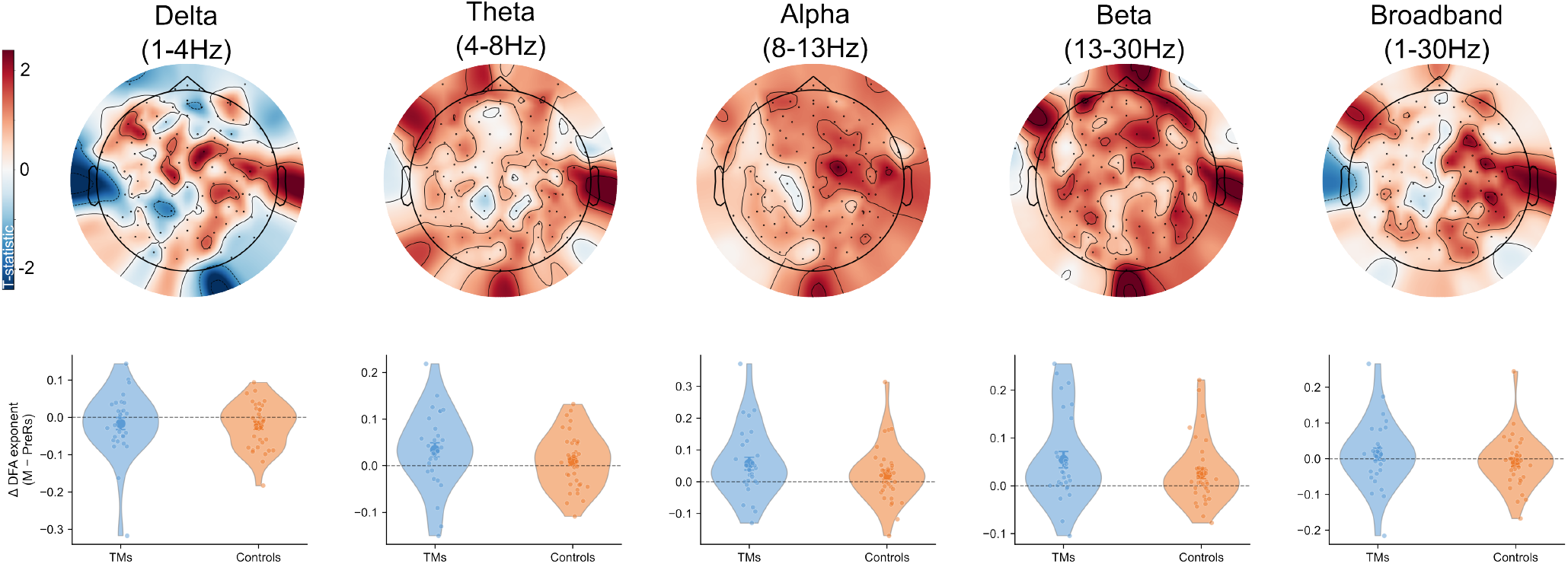
Differences in DFA exponents between groups and across frequency bands for the TM/counting condition.. **(Top)** T-values of the baseline-corrected difference in DFA exponents between TMs and controls conducting either TM or the control counting condition. No cluster-based permutation tests reached significance after FDR correction. **(Bottom)** Whole-brain baseline-corrected values for TMs (blue) and controls (orange) during the TM/counting conditions.

**Figure S3:**
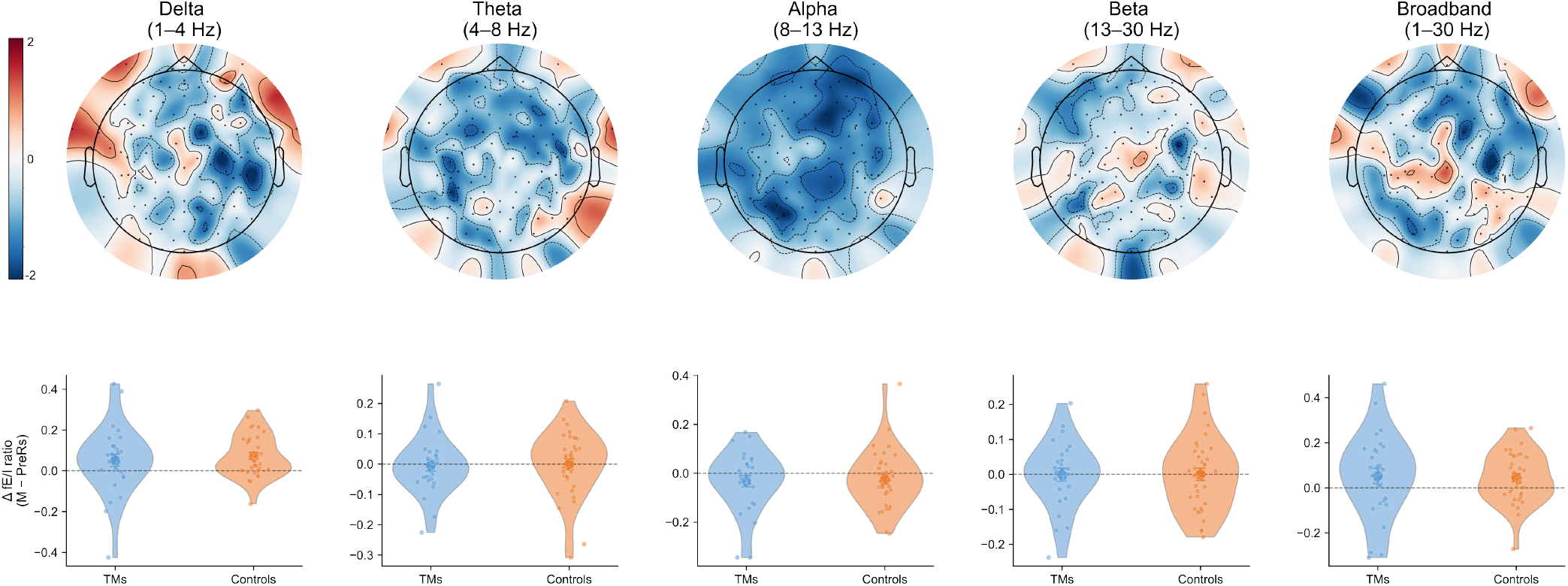
Differences in fEI values between groups and across frequency bands for the TM/counting condition.. **(Top)** T-values of the baseline-corrected difference in fEI values between TMs and controls conducting either TM or the control counting condition. No cluster-based permutation tests reached significance after FDR correction. **(Bottom)** Whole-brain baseline-corrected values for the corresponding frequency band (column) for TMs (blue) and controls (orange) during the TM/counting conditions.

**Table S1:**
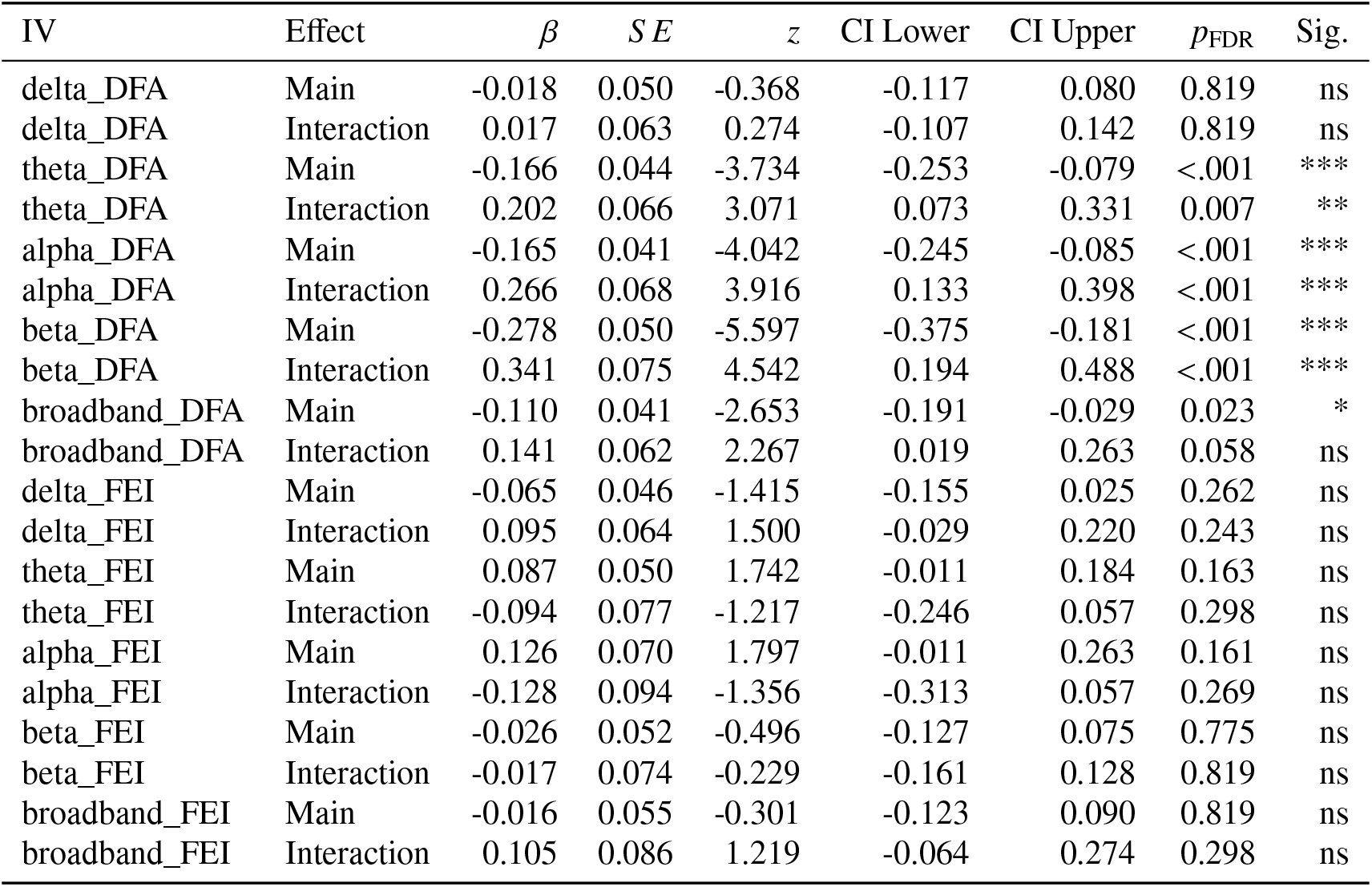
Mixed-effects model results for each criticality metric predicting TET subjective effort. Sig. is based on *p*_FDR_; ns = *p* > 0.05, * = *p* < 0.05, ** = *p* < 0.01, *** = *p* < 0.001.

| IV | Effect | $\beta$ | $SE$ | $z$ | CI Lower | CI Upper | $p_{\text{FDR}}$ | Sig. |
| --- | --- | --- | --- | --- | --- | --- | --- | --- |
| delta_DFA | Main | -0.018 | 0.050 | -0.368 | -0.117 | 0.080 | 0.819 | ns |
| delta_DFA | Interaction | 0.017 | 0.063 | 0.274 | -0.107 | 0.142 | 0.819 | ns |
| theta_DFA | Main | -0.166 | 0.044 | -3.734 | -0.253 | -0.079 | <.001 | *** |
| theta_DFA | Interaction | 0.202 | 0.066 | 3.071 | 0.073 | 0.331 | 0.007 | ** |
| alpha_DFA | Main | -0.165 | 0.041 | -4.042 | -0.245 | -0.085 | <.001 | *** |
| alpha_DFA | Interaction | 0.266 | 0.068 | 3.916 | 0.133 | 0.398 | <.001 | *** |
| beta_DFA | Main | -0.278 | 0.050 | -5.597 | -0.375 | -0.181 | <.001 | *** |
| beta_DFA | Interaction | 0.341 | 0.075 | 4.542 | 0.194 | 0.488 | <.001 | *** |
| broadband_DFA | Main | -0.110 | 0.041 | -2.653 | -0.191 | -0.029 | 0.023 | * |
| broadband_DFA | Interaction | 0.141 | 0.062 | 2.267 | 0.019 | 0.263 | 0.058 | ns |
| delta_FEI | Main | -0.065 | 0.046 | -1.415 | -0.155 | 0.025 | 0.262 | ns |
| delta_FEI | Interaction | 0.095 | 0.064 | 1.500 | -0.029 | 0.220 | 0.243 | ns |
| theta_FEI | Main | 0.087 | 0.050 | 1.742 | -0.011 | 0.184 | 0.163 | ns |
| theta_FEI | Interaction | -0.094 | 0.077 | -1.217 | -0.246 | 0.057 | 0.298 | ns |
| alpha_FEI | Main | 0.126 | 0.070 | 1.797 | -0.011 | 0.263 | 0.161 | ns |
| alpha_FEI | Interaction | -0.128 | 0.094 | -1.356 | -0.313 | 0.057 | 0.269 | ns |
| beta_FEI | Main | -0.026 | 0.052 | -0.496 | -0.127 | 0.075 | 0.775 | ns |
| beta_FEI | Interaction | -0.017 | 0.074 | -0.229 | -0.161 | 0.128 | 0.819 | ns |
| broadband_FEI | Main | -0.016 | 0.055 | -0.301 | -0.123 | 0.090 | 0.819 | ns |
| broadband_FEI | Interaction | 0.105 | 0.086 | 1.219 | -0.064 | 0.274 | 0.298 | ns |

## Footnotes

1 The authors interpret their results as a shift towards critical dynamics, through the modulation of both sensory perturbation and increased neural signal diversity.

2 Subjective effort was baseline-corrected so that the first value of each participant’s TET trace was 0, and the succeeding values were relative to this first value.

3 Available values of effort refer to stretches of data that had no epochs removed, and could therefore be used within the neurophenomenological analysis. This does not necessarily mean that the periods used in the analysis were the lowest subjective effort for the whole 30-minute meditation.

## Notes

### Competing Interest Statement

The authors have declared no competing interest.

